# *In situ* classification of cell types in human kidney tissue using 3D nuclear staining

**DOI:** 10.1101/2020.06.24.167726

**Authors:** Andre Woloshuk, Suraj Khochare, Aljohara Fahad Almulhim, Andrew McNutt, Dawson Dean, Daria Barwinska, Michael Ferkowicz, Michael T. Eadon, Katherine J. Kelly, Kenneth W. Dunn, Mohammad A. Hasan, Tarek M. El-Achkar, Seth Winfree

## Abstract

To understand the physiology and pathology of disease, capturing the heterogeneity of cell types within their tissue environment is fundamental. In such an endeavor, the human kidney presents a formidable challenge because its complex organizational structure is tightly linked to key physiological functions. Advances in imaging-based cell classification may be limited by the need to incorporate specific markers that can link classification to function. Multiplex imaging can mitigate these limitations, but requires cumulative incorporation of markers, which may lead to tissue exhaustion. Furthermore, the application of such strategies in large scale 3-dimensional (3D) imaging is challenging. Here, we propose that 3D nuclear signatures from a DNA stain, DAPI, which could be incorporated in most experimental imaging, can be used for classifying cells in intact human kidney tissue. We developed an unsupervised approach that uses 3D tissue cytometry to generate a large training dataset of nuclei images (NephNuc), where each nucleus is associated with a cell type label. We then devised various supervised machine learning approaches for kidney cell classification and demonstrated that a deep learning approach outperforms classical machine learning or shape-based classifiers. Specifically, a custom 3D convolutional neural network (NephNet3D) trained on nuclei image volumes achieved a balanced accuracy of 80.26%. Importantly, integrating NephNet3D classification with tissue cytometry allowed *in situ* visualization of cell type classifications in kidney tissue. In conclusion, we present a tissue cytometry and deep learning approach for *in situ* classification of cell types in human kidney tissue using only a DNA stain. This methodology is generalizable to other tissues and has potential advantages on tissue economy and non-exhaustive classification of different cell types.

## Introduction

Tissue organs in the human body are made up of cells that are highly specialized and organized into functionally important architectures. Identifying the various types of cells and their spatial organization within the tissue is essential to understand the physiological functions of organs and their dysfunction during disease. For example, the organization of kidney tissue illustrates how distinctly heterogeneous cell types work harmoniously to achieve blood filtration and homeostasis and how alteration in function can induce pathological states during kidney disease.^1–4^ Although novel technologies such as single cell RNA sequencing provide a new development to classify cell types and subtypes based on transcriptome profiling in disaggregated tissues,^5–7^ the ability to classify various cell types *in situ* based on imaging data, particularly in the human kidney, is not yet fully developed. The development of imaging analytics including classification techniques is crucial, as preserving the tissue architecture and spatial context of each cell will enhance the ability to interpret how specific cell types are linked to biological function.

Cell identification in tissue specimens can be performed in thin sections using histological stains such as Hematoxylin and Eosin (H&E), or Period acid-Schiff (PAS). These stains label the nuclei of cells distinctly from the cytoplasm, thereby allowing for cell visualization and detection. The identification of cell types in such imaged sections typically depends on the expert eye of a pathologist, although newer computer-aided decision support and machine learning tools have enabled enhanced reproducibility. ^8^,^9^ The use of such tools in cell classification has predominantly focused on detection of cancer or unique cell phenotypes.^10^,^11^ Broader application of these machine learning approaches in cell classification requires large training datasets, which are difficult to generate. ^9^ In this setting, it is also difficult to classify cells into multiple subtypes that could be linked to biological functions without the use of specific markers. Therefore, to enhance the feasibility and depth of classification, concurrent staining for multiple markers that are individually unique for specific cell populations is needed. Although such multiplexed staining is possible using immunohistochemistry, this approach is typically done using immunofluorescence microscopy because of the availability of established methods of quantitating fluorescence and the ability to multiplex spectrally distinguishable fluorophores for various cell markers in a single section of tissue.^12–14^ With optical sectioning microscopy, querying cell types based on specific markers could also be expanded to 3-dimensional (3D) space, increasing the ability to assay cellular structure and tissue architecture.^13^,^15^,^16^

Multiplex imaging based on cell markers may offer a path to classify cells in tissues, especially with the availability of tissue cytometry software tools that facilitate the analytical process. For example, we recently described the Volumetric Tissue Exploration and Analysis (VTEA) tool which enables the semi-automated classification of labeled cells in 3D image volumes.^13^,^15^,^17^ However, multiplexed imaging and tissue cytometry have several challenges that may limit their utility for comprehensive cell classification *in situ*. These challenges include a potentially lengthy technical workflow, a requirement for image processing expertise to optimize the analysis, and above all, only a finite number of cell-associated markers can be obtained from a single experiment.^15^ The scarcity of markers is the most limiting, as each time a new marker is discovered or needed, new experiments on additional tissue sections are required. For sparse tissue such as a kidney biopsy, this could lead to tissue exhaustion. Furthermore, because multiplexing is performed at the experimental level, classification of cells in historical datasets from sparse tissue specimen using prospectively discovered new markers is not feasible using this standard workflow. Therefore, it is beneficial to classify cells independently of the presence of specific cell markers in each experimental condition. Relying on information from a common cell marker easily embedded in the staining process and biologically linked to different cell types in the tissue could substitute the need for specific cell markers.

In this work, we proposed a hypothesis that 3D nuclear staining with 4’,6-diamidino-2-phenylindole (DAPI), a nuclear stain commonly used in most fluorescence imaging methods, ^18^ contains enough information for reliable classification of human kidney cells *in situ* using a supervised learning framework. This hypothesis is supported by previous *in vitro* work from multiple investigators showing that nuclear staining can infer functional information about cells.^9,19–21^ This is not surprising because healthy cell functions (such as various stages of the cell cycle and gene expression) and injury states are associated with specific patterns of chromatin condensation. ^21^,^22^ Furthermore, different cell types may have different shapes of nuclei, which is also captured by nuclear staining.^13^ To test this hypothesis, we leveraged an enhanced functionality of the VTEA cytometry tool to generate large ground truth datasets from kidney tissue labeled with specific markers and investigated the accuracy of various classification methods.

Our experimental results show that deep learning (DL) based classification models are able to perform kidney cell classification with a satisfactory accuracy, outperforming other classical supervised classification approaches, and that nuclear morphology can successfully classify most cell types in the cortex of the kidney by DL. Furthermore, our results suggest that 3D data improves the predictive ability of this approach compared to 2D data. In addition, minimal labeled data is needed to classify new specimens, which broadens the use of this technique for fine tuning new experiments. The approach of combining tissue cytometry (VTEA) and DL to generate ground truth library datasets as well as classify and visualize cell types *in situ* is unique. We anticipate that our methodologies will have an important impact to facilitate comprehensive cell classification and visualization *in situ* within the kidney and potentially other tissues and may enhance the information obtained from sparsely available human specimens. The datasets and approaches used will be made publicly available.

## Results

### Nuclear morphology and staining pattern are unique to distinct cell types found in the cortex of the human kidney

Within the kidney, nephron structures along with the surrounding blood vessels and interstitium are divided into specific segments based on their spatial location and physiological function ^4^ (**Figure 1A**). These segments are comprised of unique and specialized cells that can be identified by specific markers,^3^,^5^,^23^,^24^ such as Megalin (LRP2) and Aquaporin 1 (AQP1) for proximal tubular cells (PT), Tamm-Horsfall protein (THP) for thick ascending limb cells (TAL), SLC12A3 for cells of distal convoluted tubules (DCT), Cytokeratin 8 (CYT8) for collecting duct cells (CD), CD31 for endothelial cells (subdivided into two subclasses based on association with glomeruli or tubules/interstitium), Nestin for podocytes (Podo), and CD45 for leukocytes. Based on these markers, these different cells types were visually identified using confocal fluorescence 3D imaging (**Figure 1C**). The DAPI nuclear stain were qualitatively examined for specific cell types and found to have noticeable distinct signatures that are imparted by various patterns of chromatin condensation and unique shapes for each cell type (**Figure 1B**).

**Figure 1.**
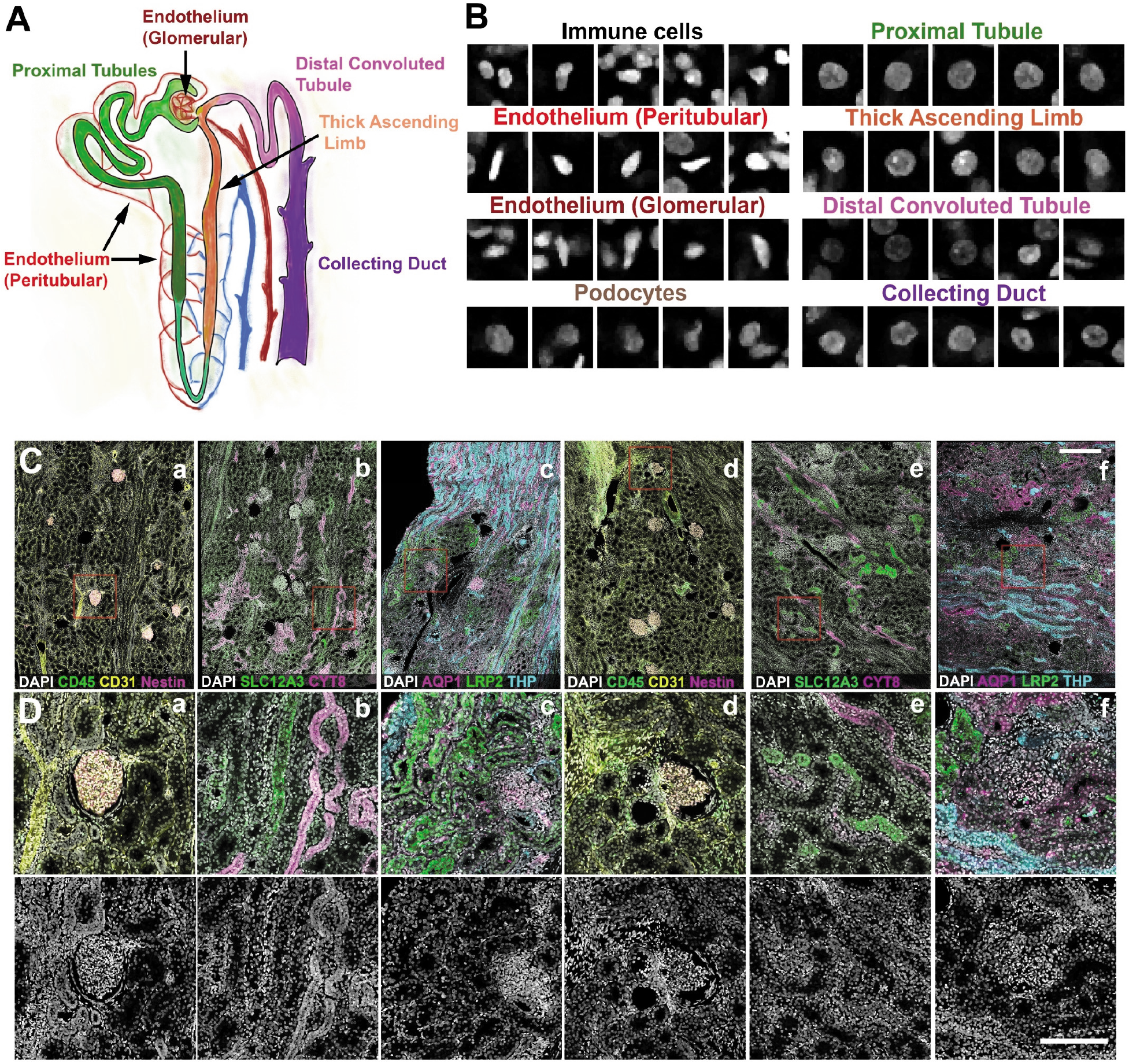
Uniqueness of nuclear morphology and staining signature of cell types found in the human kidney. **A**. Unique tubules and structures found in the nephron unit of the kidney. **B**. DAPI nuclear staining signature of various cell types identified manually based on specific markers (from C), location and morphology. **C.** 50 μm sections from human nephrectomy specimens were stained with three sets of markers all including DAPI in three different experiments. Tissue was imaged by tile scanning confocal microscopy and the images stitched together. **D**. Subregions from indicated red boxes in C. Bottom panels showing the DAPI channel only, indicate the number and variety of nuclear morphology present in the cortex of the human kidney. Scale bars = 200 μm.

### Tissue cytometry using the Volumetric Tissue Exploration and Analysis (VTEA) software tool can be used to generate large amounts of training data with cell type labels

VTEA cytometry tool was used to identify all cells based on a previously described nuclear segmentation and cytometry approach.^13^ Cell types were identified based on the fluorescence intensity of specific cell markers using an unsupervised machine learning clustering algorithm (**Figure 2**). Visualization of the specific clusters in the image volume was also done using VTEA as an additional measure to validate the identity of cell types based on the expected spatial distribution and morphological characteristics in the tissue (**Figure 2**). Image regions-of-interest were also used to limit specific localization dependent sub-populations, such as the endothelial cells in glomerulus vs peritubular space. VTEA supports projections and export of 3D volumes that can include surrounding signal or only the nuclei segmented by VTEA (**Figure S1**). Using this approach, ~230,000 cells were classified into 8 different classes (**Table S1**). The corresponding 3D volumes, 3D volumes with context (**Figure S1B**) and 2D projections along the z-axis (z-projections) were sorted and exported into “ground truth” libraries as NephNuc3D, NephNuc3D with context and NephNuc2D_Projection (without or with context), respectively (entire dataset will be available through an online repository such as the Broad Bioimaging Benchmark Collection)^25^. The median z-axis slices of nuclei were extracted from the 3D data to generate 2D datasets (NephNuc2D, without or with context).

**Figure 2.**
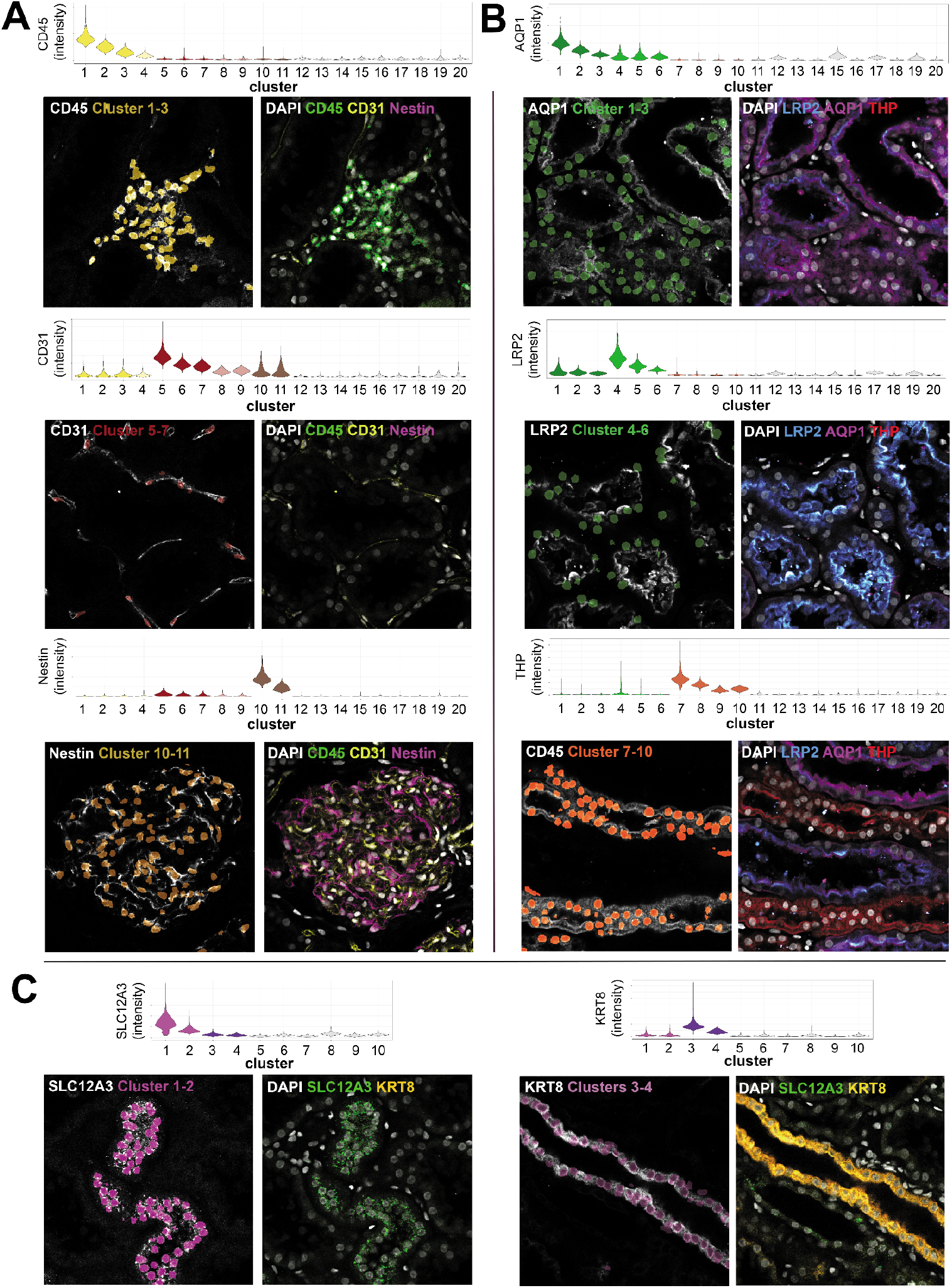
Training set generation and validation of cell type images. 50 μm sections from human nephrectomy specimens were stained with three sets of markers all including DAPI in three different experiments. Tissue was imaged by tile scanning confocal microscopy and images stitched together and processed for tissue cytometry by VTEA. Cells were classified by X-means clustering based on their associated marker intensity by unsupervised machine learning as outlined in the methods. Classified cells were mapped by cluster color on violin plots. Mapping of identified clusters is displayed on the left of each panel and original volumes at shown at the right. Tissue sections were stained with CD45, CD31 and Nestin (**A**), AQP1, LRP2 and THP(**B**) and SLC12A3 and KRT8 (**C**).

**Figure 3.**
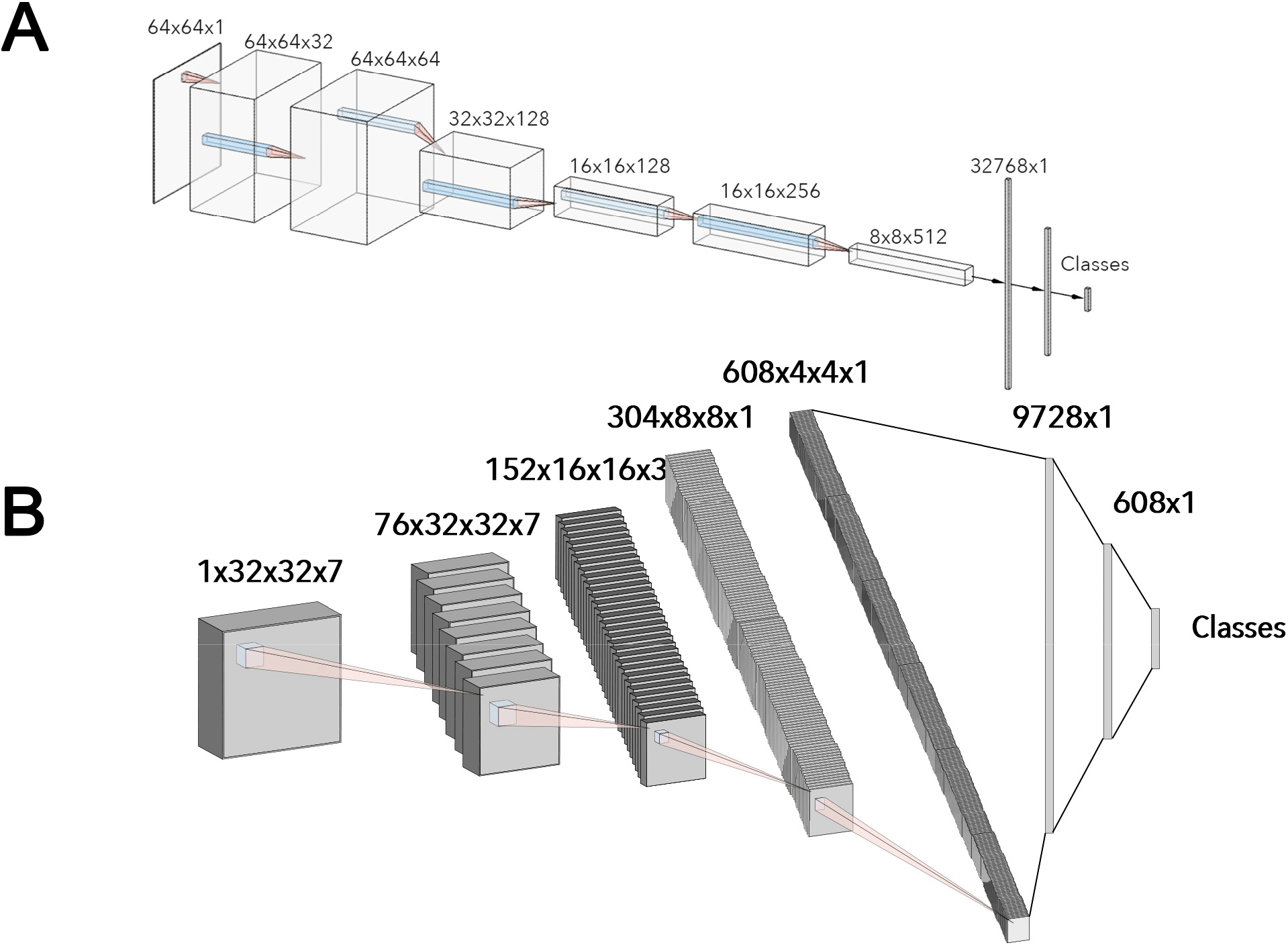
CNN Architectures for NephNet2D and NephNet3D. CNNs were developed and implemented in PyTorch. **A.** 2D CNN architecture, where each layer is separated by one 3×3 convolution, batch normalization, leaky ReLU, and max pooling with a stride of 2×2×2. Linear layers are separated by Dropout normalization (p=0.5). **B.** 3D architecture where each layer consisted of two 3×3×3 convolutions, batch normalization, leaky ReLU, and max pooling with a stride of 2×2×2. Linear layers are separated by dropout normalization (p=0.5).

### Using classical machine learning and shape-based descriptors for classifying cells based on DAPI nuclear staining

The training dataset generated was used to test the accuracy of traditional machine learning tools, including Random Forest or Naïve Bayes to classify cell types based on the DAPI 3D nuclear staining (**Figure S2, Table S2**). Three distinct models were each trained using a different feature-sets. The first feature-set used NephNuc3D data, the second feature-set considered NephNuc3D with context, and the final feature-set used NephNuc2D_Projection without context, in which the largest pixel intensity of the z-axis was used. The balanced accuracies of a Random Forest classifier for the three feature-sets were 35.0%, 33.2% and 32.7%, respectively (**Figure S2B, Table S2**). The performance of a Naïve Bayes classifier was even lower than Random Forest for all three feature-sets (**Figure S2, Table S2**).

Similarly, we also tested the accuracy of a shape-based descriptor, using spherical harmonics (SPHARM), in predicting cell classes.^26^ SPHARM analysis was performed after surface extraction using the marching cube algorithm.^27^ Once the spherical harmonic features of the images were obtained, both Support Machine Vector and Random Forest classifiers were used as classification models (**Figure S3, Table S2**). The best-balanced accuracy using SPHARM was 15.5% with the NephNuc3D data.

### Convolutional neural network (CNN) based Deep Learning models increase the accuracy of cell classification

Next, a deep learning approach was tested to determine if we could improve the classification accuracy. For this, two convolutional neural networks, NephNet2D and NephNet3D were devised to test the 2D or 3D training data, respectively. The input for NephNet2D were NephNuc2D_Projection data or median z-axis slice 2D datasets. The input for NephNet3D were NephNuc3D or NephNuc3D with context. The architectures of the networks used are shown in **Figure 2**. Augmentation and hyperparameters selection settings are discussed in detail in the Methods section. The inclusion or exclusion of contextual, surrounding nuclei in the 2D or 3D data was examined to improve the accuracy.

Both the dimension and content of the images affected the predictive ability of the CNNs. For the 2D slice, 2D maximum projections, and 3D volumes, adding the context surrounding the nuclei of interest improved the balanced accuracy (**Figure 4** and **Table S2)**. The highest balanced accuracy was observed with NephNet3D using NephNuc3D with context (80.3%), as compared to 66.5% and 60.8% with NephNet2D using NephNuc2D_Projection and NephNuc2D with context, respectively. Surprisingly, using an established vision-trained network such as the fine-tuned Resnet-31 CNN underperformed compared to NephNet2D (**Figure S4 and Table S2**).

**Figure 4.**
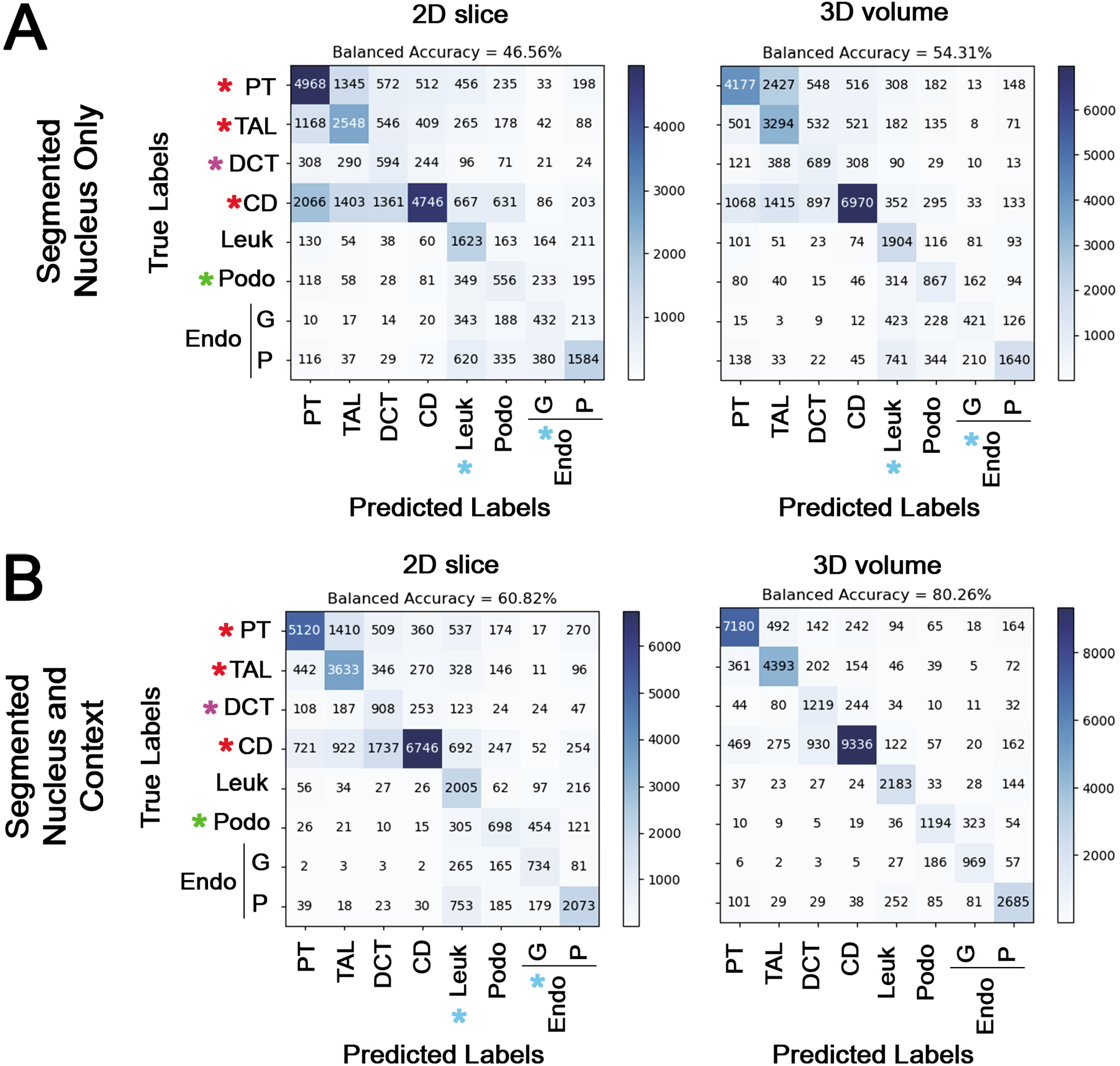
Cell classification based on nuclear staining using NephNet2D or NephNet3D and the NephNuc datasets. The NephNuc datasets were split into training and testing. The eight classes used for training are epithelial cells from the proximal tubules (PT), thick ascending limbs (TAL), distal convoluted tubules (DCT), and collecting duct (CD), and other cells such as leukocytes (Leuk), podocytes (Podo) and endothelial cells (Endo) in glomeruli (G) or in the peritubular (P) space. The testing datasets were classified, and accuracy and confusion matrices were generated. **A.** The balanced accuracies of networks trained on 2D sections (left) or 3D volumes (right) containing a single nucleus. **B**. The balanced accuracies of networks trained on 2D sections (left) or 3D (right) containing a nucleus and surrounding nuclei. Asterisks indicate specific weaknesses in either the 2D or 3D classifications and the influence of surrounding nuclei on the classification. In all configurations except 3D nuclei with surrounding nuclei, there were errors in classifying podocytes as leukocytes and glomerular endothelium (blue and green asterisks) and between epithelial cells (red asterisks). Surrounding nuclei, context, improved classification of DCT (magenta asterisks, compare A to B).

The network with the highest balanced accuracy, NephNet3D trained on 3D image volumes with context, was subjected to noise and image resolution robustness testing (**Figure 5**). Down sampling the input images led to a rapid decrease in predictive performance. For example, a 2x down sample decreased the accuracy from 80.26% to 65.49%. The addition of noise at fixed levels mildly decreased the predictive performance, dropping from 80.26% to 77.45% at an α = 0.8 and 71.39% at an α = 0.6.

**Figure 5.**
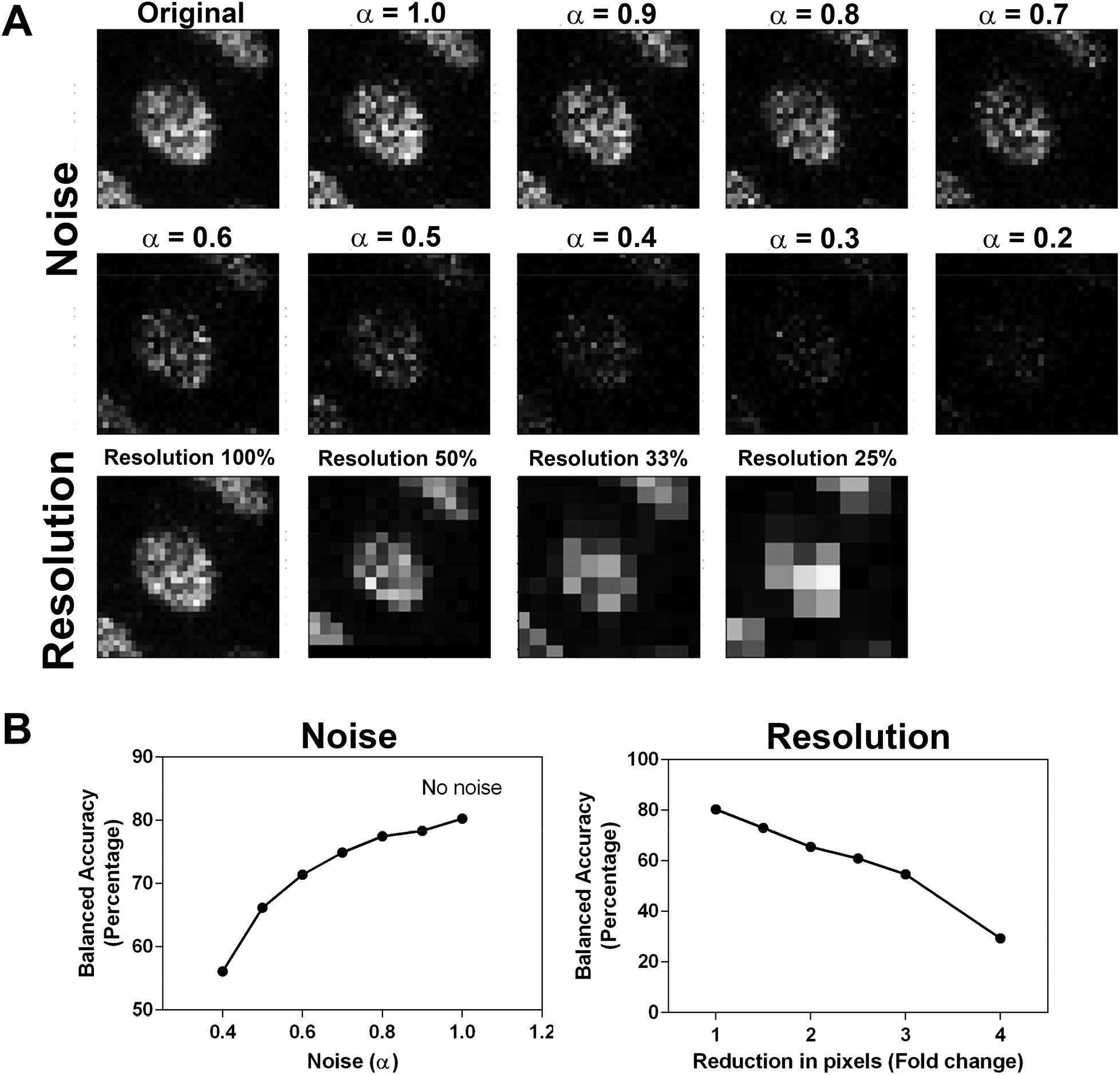
NephNet3D performance on noisy and lower resolution images. Increasing amounts of noise or decreasing resolution was used to generate testing datasets of 3D nuclei from the NephNuc3D with context data and classified with NephNet3D. A nearest neighbor approach was used for reducing the resolution of the image and noise was added by incorporating an α-factor described in the methods. **A.** Example of augmentation of training data with increasing noise or decreasing resolution for one nucleus. **B**. NephNet3D performance on testing data augmented by adding noise (left) or reducing the number of pixels to simulate less resolution (right).

### Improving classification in novel specimens’ accuracy by subsampling

To improve the predictive ability of NephNet3D, multiple samples were included in the NephNuc datasets. As specimens from novel patients are analyzed, we expect there will be specimen-to-specimen variability. Thus, during training of NephNet3D, we tested if the ability to predict cell classification in a new specimen was improved by fine-tuning on fractions of that specimen. An early iteration of NephNet3D (trained only on specimens 1 and 2(**Table S1**)) was trained for 30 epochs on 1% of specimen 3. This finetuning improved the balanced accuracy on the remaining nuclei of specimen 3 from 58.3% to 74.2%. With 10% of specimen 3 used for fine-tuning, the network’s balanced accuracy was 79.7%-nearly the same balanced accuracy of the fully trained NephNet3D network. When finetuning with more than 10% of specimen 3, the balanced accuracy remained at ~80%. This suggests that, if necessary, NephNet3D can be adapted to novel tissue with a small fraction of labeled nuclei (**Figure 6A**).

**Figure 6.**
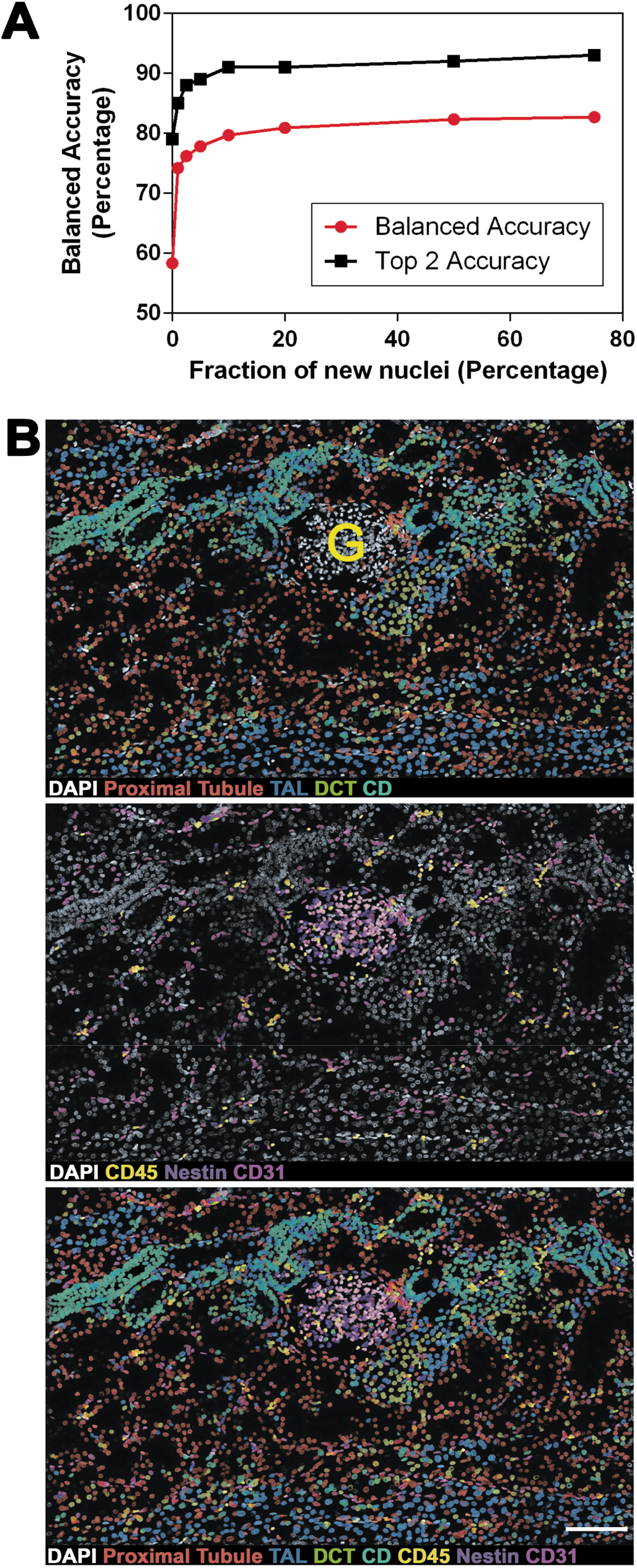
NephNet3D classification of cells in new image volumes. **A.** Finetuning of NephNet3D with nuclei from specimen 3 labeled for the eight cell types. 10 percent of specimen 3 for fine tuning of NephNet3D trained on specimens 1 and 2 is sufficient for near-peak balanced accuracy on the entire specimen. The CNN was fine-tuned on varying amounts of data from a new specimen (nephrectomy) prior to being tested on the nuclei from the new specimen. **B.** An image volume of DAPI stained nuclei not previously used, from specimen 1, was segmented by VTEA and classified with NephNet3D. Overlay of predicted labels from NephNet3D on the DAPI stained nuclei. The overlays in the top panel are for cells classified as: proximal tubules (red), TAL (blue), DCT (yellow-green) and CD (blue-green); in the middle panel, leukocytes (CD45, yellow), podocytes (Nestin, purple) and endothelium (CD31, magenta and pink). Bottom panel shows DAPI stained nuclei with all predicted labels. G indicates a glomerulus. 177 segmented object/nuclei were manually classified by an expert. NephNet3D had an agreement of **67.9%** (balanced accuracy as compared to expert classified nuclei). Maximum Z-projections are shown, scale bar = 100 μm.

### Improving classification by training data augmentation

Supervised classification generally performs better if the training data has sufficient variability. This is particularly true for deep learning-based classification, as such models have very large number of parameters; unless the training data has sufficient variability, the large number of parameters causes the model to succumb to overfitting. To overcome this problem, data augmentation is used during training to increase data variability and mitigate the overfitting issue. The image augmentation steps we considered included transformation, rotation, flip and noise injection. The balanced accuracy on training without augmentation were 13%, 45%, and 53% for 2D model, 3D model without context, and 3D model with context, respectively. In contrast, training using augmented training data yielded a balanced accuracy of 29%, 53%, and 80% for 2D model, 3D model without context, and 3D model with context data, respectively.

### NephNet3D with VTEA tissue cytometry can classify and spatially map cell types in image volumes stained only with DAPI

The ultimate goal of this methodology is to classify cell types in image volumes based only on the DAPI nuclear staining. To establish feasibility, a new image volume of a kidney cortical section stained with DAPI that was not used in previous training and from the same specimen cohort was tested. Image volumes of all the nuclei were generated by VTEA and then tested through the trained NephNet3D to predict the classes of the cells. Classified nuclei were then visualized on the image volume using VTEA, which allowed a “pseudo” highlighting of cells based on the trained NephNet3D (**Figure 6B**). These predictions by NephNet3D were compared to classification by an expert (using nuclear shape and spatial cues) on a subsample of the image volume. Across all eight classes, NephNet3D had a balanced accuracy of 67.9% as compared to an expert’s classification. The NephNet3D and expert classifications showed fair agreement with a Kappa statistic of 0.35. Furthermore, upon examining the cell classifications mapped back to the image volume, cells classified as epithelial cells outlined contiguous tubular structures (**Figure 6B, top panel**) while cells classified as endothelial or immune cells were confined to either the interstitium or glomeruli (**Figure 6B, bottom panel**). Strikingly, cells classified as podocytes were almost exclusively confined to the single glomerulus in the image volume (**Figure 6B**).

## Discussion

In this work, we devised an approach to classify cells *in situ* within intact kidney tissue, using an imaging-based approach that relies on nuclear staining. To accomplish this goal, we used tissue cytometry to efficiently generate a large amount of training data, consisting of image volumes of nuclei classified based on their association with specific cell markers within the human kidney. This dataset was used to determine the optimal machine learning approach that could provide the highest classification accuracy based on this nuclear staining. Our results show that a deep learning approach outperforms traditional supervised classification methods, and that a CNN trained on 3D image volumes with context (other nuclei in vicinity) provides the highest balanced accuracy. This classification pipeline was then demonstrated on an unexplored kidney tissue whereby cells were successfully classified into 8 subtypes and visualized based only on 3D nuclear staining.

Image-based classification of cells has several applications in biology and medicine.^9^ Some commonly recognized applications are in drug screening and discovery,^19^,^28^ genetic screening, ^29^ cell biology,^30^ and digital pathology. ^10^ These last two application are commonly used in the context of cell classification in intact tissue. Cell biology applications include cell classification based on nuclear and other specific markers or spatial analyses, similar to what we and other described as tissue cytometry using fluorescence or histochemical markers. ^12^,^13^,^16^,^31^,^32^ Many efforts are also focused on cell segmentation, automated detection and counting.^33^ Digital pathology relies on histology staining and has experienced exciting development in cell segmentation and classification. ^8^,^34–36^ Several approaches have been described to classify cells based only on nuclear staining in digital pathology images, such as standard machine learning classifiers, numerical feature engineering, neural networks and transport-based morphometry. ^9^,^10^,^20^ However, many of these approaches are focused on cancer detection or identification of unique cell phenotypes. ^10^,^11^ An approach that can allow non-exhaustive cell classification based only on 3D nuclear signature has not been previously described.

A strength of our approach is that it uniquely combines tissue cytometry and deep learning to classify cells based on their 3D nuclear staining in intact tissues. Tissue cytometry using VTEA is essential to generate the training dataset and visualize the classified cell *in situ*. Generating training data is arguably one of the most arduous tasks in any machine learning approach. VTEA allowed us to generate 230,000 labeled nuclei quickly-in a matter of weeks. Furthermore, VTEA will keep track of the nuclei and map them to the tissue for visualization after classification, allowing their classification to be interpreted in the setting of their spatial distribution and relationship to other cells and structures. In the kidney, this is critical in interpreting how specific cell types are linked to biological function or dysfunction, and we envision that this will also be important when applied to other organs.

The approach proposed in this work will likely have important applications in the setting of sparse tissue. The described workflow enables continued re-evaluation of existing imaging data, thereby reducing the need for additional tissue. Indeed, as new cell markers emerge, the deep learning approach can be trained to recognize these new nuclear signatures in ground truth datasets generated in an abundant source of tissue, such as the specimen used in this study. New classifications or sub-classifications can then be imputed on existing datasets from sparse tissue without the need for additional sectioning and depletion of the tissue. This is especially useful in the setting of a kidney biopsy, which is a needle biopsy with a small amount of tissue.^17^,^37^ DAPI staining can be easily incorporated with other technologies. For example, our approach can be extended to specimen prepared for RNA exploration, and cell classification can become highly integrated by incorporating signatures from various levels of gene expression. ^38^ Although this overall approach has been tailored to the complexity of kidney tissue, it can be applied to other organs. Furthermore, the model of continuous data extraction from a single archived dataset may facilitate data sharing and collaboration.

Comparing the different approaches and networks support the conclusion that a 3D CNN architecture was best at predicting cell class based on nuclear staining and showed marked improvement over a 2D architecture tested with various types of data. The 3D CNN, with 3D data, was particularly important for improving the classification accuracy of cells within the glomerulus and proximal tubules. Furthermore, NephNet3D showed increased accuracy for the podocyte, proximal tubule, thick ascending limb, collecting duct, and glomerular endothelium cell classes. This improvement supports the idea that important features are contained within the three-dimensional structure of the nucleus for each cell type. Importantly, NephNet3D did have trouble classifying or discriminating between tubular epithelium of adjacent segments, such as the collecting duct and distal tubule. This could be due to the biological reliability of the cell markers, the variability in expression of the markers (e.g. Sample 3, Table S1) or overlap in transition areas between tubular segments ^23^. In fact, the poor predictive capacity in these transitional regions (e.g. DCT to CD) could be used as a method to identify transitional cells or possibly novel subtypes of the tubular epithelium.

The wide availability of 2D imaging modalities, clinically and in research, argues for developing 2D approaches for cellular classification. We demonstrate fair performance (~60% balanced accuracy) of a 2D architecture. We also demonstrate poor performance (~30% balanced accuracy), worse than our 2D approach, of a pretrained network, Resnet-31^39^. Thus, our 2D network could be a better starting point for additional refinement of a 2D approach. Furthermore, our work demonstrates the importance of data collection at an appropriate resolution and signal/noise for the predictive ability of deep learning using nuclear information. This may provide a guide for future data collection on additional imaging modalities more common in 2D imaging, including slide-scanners and other high-content platforms. Importantly, we expect both our 2D and 3D architectures will be refined through additional data collection to establish more classes of nuclei, more depth within classes and to cover other imaging platforms and modalities. Our network and data are publicly available to the community for such development.

Our study has several limitations. The approach used is specialized and may not be broadly applicable because of the need for expertise in imaging and computer science as well as the need for high performance computing hardware. Despite having a training dataset of thousands of curated nuclear volumes, we anticipate that the training dataset will need to expand significantly for the accuracy to improve. Since the dataset has been generated from just three tissue samples, expanding the dataset with additional specimens will further improve the generalizability of this approach. Also, the resolution at which the data was collected, and subsequent quality of segmentation, may place SPHARM or numerical feature-based approaches at a disadvantage *a priori*, because these approaches may require a higher resolution image to be most effective. However, balancing feasibility and the economy of time/resource constraints, the current data acquisition settings are commonly used.^15^ Therefore, it is reasonable to assert that for standard large scale 3D image acquisition, NephNet3D is likely the most optimal approach for cell classification based on nuclear staining.

The current work was done on reference tissue from donor nephrectomy specimens. Kidney disease will likely alter the nuclear staining signature of cells because of metabolic stress or cellular injury, ^40^ thereby potentially changing the classification of some cells or presenting a new subclass for a particular cell type.^21^ Investigating these changes and distribution at the tissue level will likely aid in understanding the pathophysiology of the disease, especially in early stages where such subtle cellular changes may precede obvious structural abnormalities. We propose that the cytometry approach will facilitate this process. The visualization feature of VTEA can map all cell types, including diseased cell populations. By comparing localization and changes to reference tissue, diseased cells can be re-classified, and could serve as ground truth for a disease subclassification. Such iterative learning models are subject of ongoing and future investigations in our lab. In addition, since we showed that only a small fraction of appropriately classified nuclei from a tissue is enough to maximize the accuracy of cell class prediction for that particular tissue, expanding our training dataset to new subclasses of cells may conform with our goals of tissue economy, even when only sparse tissue is available for ground truth generation.

Our work here does not address which specific features in the nuclei are key determinants of their classification. The explainability and human readability of the deep learning approaches are still at an early stage.^41^ However, since the signatures of nuclear staining are linked to chromatin condensation and cellular activity,^18^,^21^ it is possible that our work could present an opportunity to recognize the salient features in nuclei that dictate their classification. For this, we plan to use class saliency map and attribution-based approaches to understand the specific image features which impacted the classification decision made by the deep learning models. This is a subject of ongoing research.

In conclusion, by combining tissue cytometry and a deep learning CNN, we present an approach for *in situ* classification of cell types in the human kidney using 3D nuclear staining. This classification methodology allows the preservation of tissue architecture and spatial context of each cell and has potential advantages on tissue economy and non-exhaustive analysis of existing data.

## Methods

### Kidney Tissue Specimens

Three deceased donor kidney nephrectomy tissue were used based on a protocol approved by the Institutional Review Board at Indiana University. 50 μm section from OCT frozen kidney cortical specimen were fixed overnight with 4% paraformaldehyde or 50 μm sections were cut by vibratome from 4% paraformaldehyde fixed cortical tissue fragments underwent staining with DAPI and antibodies for the specified markers including: Megalin, LRP2 (cat#ab76969, abcam), Aquaporin-1 (AQP1,clone 1/22, cat# sc-32737, Santa Cruz Biotechnology), Slc12A3 (cat#HPA028748, Sigma), Uromodulin, Tamm-Horsfall protein (cat# AF5144, R&D Systems), CD45 (clone HI30,cat# 304002,), CD31 (clone JC/70A cat# NB600-562-R, Novus), Nestin (clone 10C2, cat# 656810, Biolegend) and Cytokeratin-8 (cat# NBP-34267, Novus). All staining was performed in 1X Phosphate buffer saline with 10% normal goat serum and 0.1% Triton X-100 with overnight incubations. All secondaries were raised in goat highly cross-adsorbed, and were labeled with Alexa488, Alexa546, Alexa555 or Alexa568, Alexa647. The CD31, CD45 and Nestin antibodies were directly conjugated with Dylight 550, Alexa488 and Alexa 647, respectively. Stained tissue was mounted under #1.5 coverslips in Prolong Glass allowed to cure for 24-48 hours before sealing with nail-polish. Mounted and sealed tissue was stored at 4C before imaging.

### Imaging

Image acquisition was performed in four separate consecutive channels using an upright Leica SP8 Confocal Microscope controlled by LAS X software (Germany). Volume stacks spanning the whole thickness of the tissue were taken using a multi-immersion 20× NA 0.75 objective with Leica immersion oil with 1.0-μm spacing and 0.5 x 0.5 μm voxels. Large scale imaging was performed using and automated stage with volumes overlapping by ~10%. Typical imaging times were 4-6 hours. Image volumes were stitched using FIJI^42^.

### VTEA Cytometry

Tissue cytometry was performed using a prerelease version of the FIJI plugin Volumetric Tissue Exploration and Analysis(VTEA)^13^ available on github (https://github.com/icbm-iupui/volumetric-tissue-exploration-analysis, that incorporates unsupervised learning approaches into a 3D tissue cytometry workflow (manuscript in preparation). *Segmentation:* Entire image volumes were imported in VTEA and the nuclei channel was pre-processed to facilitate segmentation by performing denoising, rolling-ball background subtraction and contrast stretching to compensate for attenuation of signal at depth. A modified form of connected components built into VTEA (LSConnect3D), that combines Otsu intensity thresholding, watershed splitting and connected component merging in 3D was used for segmentation. To facilitate segmentation of mesoscale images LSConnect3D subdivides the image into user defined volumes to facilitate parallelization during processing. Both a minimum and maximum size restriction was placed on nuclei to mitigate segmentation errors. Following segmentation of the nuclei a second segmented volume, grow-volume, was defined around each nucleus that extend ~2 pixels for assessing stains associated with a nucleus^13^.The pixel values, including mean, upper-quartile mean, standard deviation and/or maximum within the segmented nuclei and/or the grow-volume were used as features for clustering by X-means implemented in SMILE (http://haifengl.github.io/).Validation of clusters was performed by an expert using VTEA’s mapping of gated cells to the original image volume to demonstrate correct localization of the selected nuclei. Nuclei from gated cells were exported by VTEA as user defined image volumes (segmented nuclei only, with surrounding nuclei or with context, image size, etc.) using a duplicate image volume of the DAPI channel that had not been subjected to the image preprocessing necessary for segmentation.

Datasets exported names and a brief description are given in table S2 and class representation are given in table S1. A small subset (<1.7%) of nuclei without context were excluded from NephNuc3D when the segmented nucleus extended outside of the sampling volume.

Image analysis was performed on a dedicated workstation with a Xeon 2245 (8 cores) with 256 GB RAM, 2×2 TB NVMe SSDs, a 24TB RAID, 2x 2080Ti (not linked, 11 GB RAM each, Nvidia) running Windows 10.

### Expert review of classification

Using the pre-release version of VTEA mentioned above, a manual classification was performed on 177 nuclei. VTEA randomly picked segmented nuclei and presented a cropped image with a highlighted nucleus for an expert to place in predefined classes. VTEA provided an interface to tally these nuclei and import these expert classifications as a feature for comparison with other classification approaches. Balanced accuracy was calculated as given below and the kappa statistic was calculated with the package *psych* in R.

### Data organization

Training data for this research consisted of approximately 230,000 3D images of nuclei, segmented as described above from three reference tissue specimens. Each grayscale image was 32×32×7 pixels representing *in situ* dimension of 17.3×17.3×3.9 μm. Each pixel value was between 0 and 255 denoting the light intensity of the pixel. Eleven kidney cell classes were identified and the ground truth class-label of each cell was obtained using VTEA cytometry tools. However, the performance of each learning model was reported by using a condensed set of eight classes; for example, while two separate proximal tubule classes are identified in the labels, classifying a nucleus into either class is recorded as a correct classification. 15% of image instances of each class were used to build a test dataset.

### Preprocessing of nuclei prior to classification

Before feeding an image to the classification algorithm, each image was pre-processed as below. First, each pixel value was normalized using Z-score normalization; if *I* was the intensity of the original pixel, *I*’ was the intensity of normalized pixel, *μ* was the mean intensity and *σ* was the standard deviation of that pixel across the training images, then *I*’ = (*I-μ*)/*σ*. This transformation made the mean intensity of the dataset to be 0 and the variance to be 1.

### Classical supervised classification models

Two classical supervised classification models were used, Random Forest, and Naïve Bayes. Random Forest classifier was implemented using Scikit-learn^43^ (version 0.22.2) in Python with a maximum number of trees equal to 450. The maximum depth was expanded until all leaves were pure or all leaves contained 2 or fewer samples. A Gaussian Naïve Bayesian classifier was used with no prior probabilities and variance smoothing factor of 1e-9.

### Classification with spherical harmonics

The method is based on the work of Medyukhina et al.^27^ Using the NephNuc3D datasets, the surface of nuclei surface was reconstructed in the form of a mesh of triangles using the marching cubes algorithm (scikit-image v. 0.18.dev0). Next, the x, y and z coordinates of unique vertices were extracted and converted to polar co-ordinates. These polar co-ordinates were used to expand a set of irregularly sampled data points into spherical harmonics using a least squares inversion (SHTOOLS 4.6), adapted from ‘https://shtools.oca.eu/shtools/public/pyshexpandlsq.html’. The maximum spherical harmonic degree of the output coefficients was set to 3. These spherical harmonic coefficients were used to compute rotation-invariant frequency spectrum to be used as static features for classification. These static features of each of the nuclei were used in a Support Vector Machine or Random-Forest classifier.

### Deep learning

A custom-made 3D deep convolution neural network (CNN) based model was used for the kidney cell classification. Besides preprocessing, which was performed prior to the training, some augmentation steps were performed during training, mainly to prevent overfitting. Each augmentation step had a 30% chance of being applied to an image on every epoch. The augmentation steps were mix of transformation, rotation, flip, and noise injection; the transformations were translation up to 8 pixels in X and Y axes and 3 pixels in Z axis, random rotation up to 35°, random 90° rotation, random horizontal or vertical flip, random Poisson noise using the equation I’ = αI_0_ + η (where I_0_ is the original pixel intensity, I’ was the transformed pixel intensity, η is the Poisson noise, and α is a mixing factor randomly chosen between 0.8 and 1.0), random down-sample up to a factor of 2x, and random contrast by multiplying the image by a random contrast factor between 0.8 and 1.2.

Three different image data formats, 2D slice, 2D maximum projection, and 3D, were considered. 2D slice (or in-short 2D) was obtained by only using the median slice of each volume. 2D maximum project was obtained by considering the maximum pixel value over the z-axis of that pixel. Each image format can include the surrounding nuclei (“with context”) or only the segmented nuclei of interest (“without context”). NephNuc3D data, the second featureset considered NephNuc3D with context, and the final feature-set used NephNuc2D_Projection without context,

For convolutional neural networks, three different architectures were considered. First, a Resnet-31 architecture was fine-tuned on the 2D or 2D maximum image data format.^44^ No weights were frozen during the fine-tuning process. Second, a CNN in which each block consists of batch normalization, 3×3×3 convolution with a stride of 2 and padding of 1, leaky ReLu, a second 3×3×3 convolution and leaky Relu, and a final maximum pooling layer with a stride of 2. These blocks were repeated until the feature size was 4×4×1 pixels. The classifier block consisted of two sets of a linear layer, batch normalization, and dropout layer (p=0.5). The initial number of features was 76 and double after each convolutional block. This architecture is referred to as NephNet3D throughout the text and received 3D images with or without context as inputs (NephNuc3D and NephNuc3D with context). Third, a similar CNN with 2D convolutions was designed such that the convolutional blocks are repeated until the feature size is 8×8 pixels. The initial number of features was 32 and doubled after each convolutional block. This architecture is referred to as NephNet2D and received 2D slices or maximum projections with or without context as input (NephNuc2D, NephNuc2D_Projection with/without context).

### Optimization of network parameters

The Hyperband optimization technique^45^was used to determine optimal values for learning rate, initial number of features, batch size, learning rate step, and learning rate decay. The Hyperband algorithm begins by selecting a predetermined number of configurations (600), and training each for 1 epoch. The top 50% determined by validation loss are trained for 2 epochs. The top 50% are selected and trained for 4 epochs. This process is repeated until only 1 network remains. Then the entire process is repeated 5 times but begins with less configurations trained for more epochs. For example, the third iteration will train 150 configurations for 4 epochs. This altered repetition strategy acts to reduce bias towards networks whose performance improves rapidly but does not achieve high performance with many additional epochs (e.g. high learning rates). The final parameters for training NephNet3D were a learning rate of 0.016, batch size of 64, momentum of 0.8, weight decay of 0.006, and 76 initial features. The learning rate schedule was optimized such that the learning rate was reduced by a factor of 0.29 after 8 epochs without an improvement in validation loss. For the NephNet2D, the learning rate was 0.000414, batch size of 16, momentum of 0.9, weight decay of 0.0058, and 32 initial features. The learning rate scheduler used a factor of 0.489 and a step size of 15 epochs without validation loss improvement.

### Training

All networks were trained at their optimized parameters for 500 epochs and tested using the network weights that achieved the lowest validation loss using PyTorch v1.5^46^. Networks were trained either on a 2080Ti (11 GB RAM, Nvidia) installed on a computer with 126 GB RAM, 6 TB of SSD space and a Core i9 9900k running Windows 10 or a workstation with two Titan RTX graphics card (24 GB RAM each) with NVLink (Nvidia), 128 GB RAM, 4 TB of SSD space and a Core i9 9900k running RedHat Linux.

#### Metrics

Due to the relatively unbalanced nature of cell types in the human kidney, a direct accuracy score would bias towards machine learning models whose majority prediction matches with the most common cell types. However, due to the importance of less common cell types, such as immune cells, a balanced accuracy was used to report the model performance. Balanced accuracy is the average of the percentage of true cells in a class correctly identified as belonging to that class, calculated as

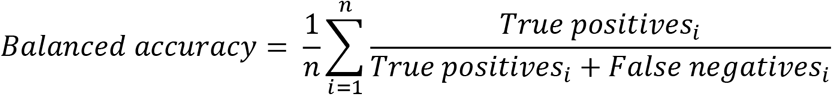

where *n* is the number of classes. This represents the average of the recall for each class and is commonly used in assessing performance on multiclass problems.

#### GitHub

The entire code base for the classical supervised classification, convolutional neural network architecture, optimization, training, and testing can be found on GitHub at https://github.com/awoloshuk/NephNet.

## Notes

### Competing Interest Statement

The authors have declared no competing interest.

