## Supplemental material for "*In situ* classification of cell types in human kidney tissue using 3D nuclear staining"

| | | Labeled nuclei per specimen (NephNuc3D) | | | |
|---|---|---|---|---|---|
| **Structure** | **Marker** | **1** | **2** | **3** | **Total** |
| Proximal tubules | LRP2, AQP1 | 16440 | 18806 | 20765 | 56011 |
| Thick ascending limb | THP | 12572 | 13031 | 9555 | 35158 |
| Distal convoluted tubule | SLC12A3 | 4944 | 6222 | 0* | 11166 |
| Collecting duct | Cytokeratin-8 | 33895 | 29236 | 12696 | 75827 |
| Immune cells | CD45 | 5099 | 11039 | 713 | 16851 |
| Endothelium (glomerular) | CD31 (spatial ROI) | 1671 | 2786 | 3927 | 8384 |
| Endothelium (peritubular) | CD31 (spatial ROI) | 1831 | 13228 | 6943 | 22002 |
| Podocytes | Nestin | 1727 | 4801 | 4483 | 11011 |

**Supplemental Table S1: Summary NephNuc3Dv1.0 dataset.** *Although the distal convoluted tubule was visible in the volume by expert identification using morphology, SLC12A3 staining proved inadequate in identifying significant numbers of nuclei.

| | | Balanced accuracy (%, by dataset) | | | | | |
|---|---|---|---|---|---|---|---|
| **Datasets** | | NephNuc 2D | NephNuc 2D | NephNuc2 D-Projection | NephNuc2 D-Projection | NephNuc 3D | NephNuc 3D |
| Approach | | *Slice without context* | *Slice with context* | *z-axis max-projection without context* | *z-axis max-projection with context* | *Volume without context* | *Volume with context* |
| *Name* | *Desc.* | | | | | | |
| Vectorized image | Vectorized images with a Random Forest classifier | 30.84 | 32.74 | ND | ND | 30.64 | 31.63 |
| Vectorized image | Vectorized images with a Gaussian Naïve Bayes classifier | 22.94 | 28.32 | ND | ND | 14.07 | 26.69 |
| SPHARM | Spherical harmonics with a Random Forest classifier | ND | ND | ND | ND | 15.53 | ND |
| SPHARM | Spherical harmonics with an SVM classifier | ND | ND | ND | ND | 12.81 | ND |
| ResNet-31 | Finetuning of existing CNN | n/a | 30.75 | 30.95 | 30.62 | ND | ND |
| NephNet2D | 2D CNN (this work) | 46.56 | 60.82 | 29.17 | 66.55 | ND | ND |
| NephNet3D | 3D CNN (this work) | ND | ND | ND | ND | 54.31 | **80.26** |

**Supplemental Table S2: Summary of accuracies for classification approaches.**

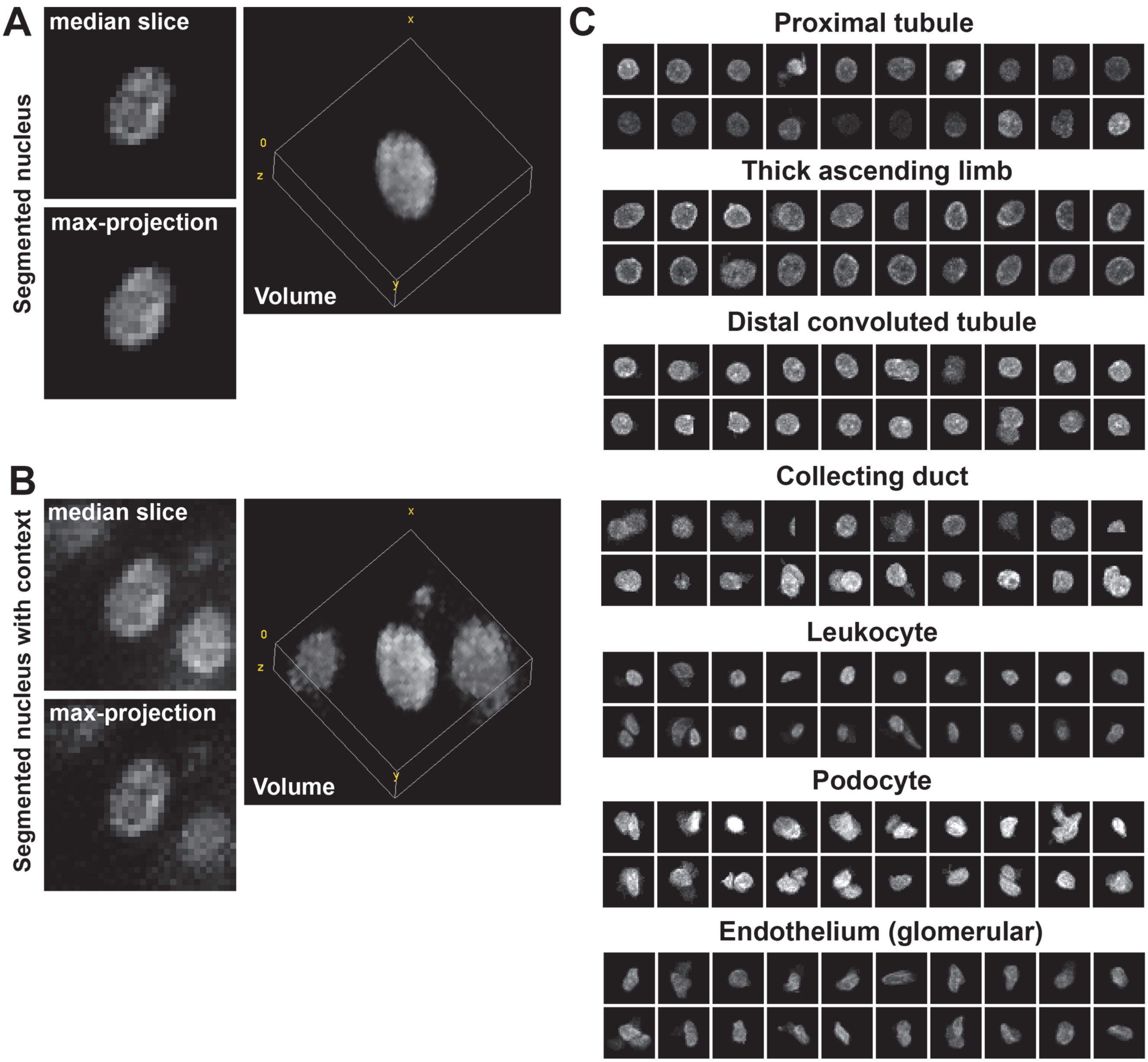


**Figure S1. Examples of training data generated by cytometry with VTEA.** Confocal mosaic image volumes of 50 µm thick sections from human nephrectomy stained with DAPI and the markers given in Figure 2. The mosaic volumes were stitched together and imported into VTEA for segmentation and classification with X-means clustering on intensity features. Images of the nuclei were extracted from the image volume. **A**. Example images of the same proximal tubule nucleus including the median, z-axis maximum projection and a volume rendering of the data used for training and testing. **B.** Example images of the same nucleus as in (A) with the surrounding nuclei used for testing the impact of context on classification accuracy. **C.** Montages of z-axis maximum projections from seven of the labels used in training and testing of classification approaches.

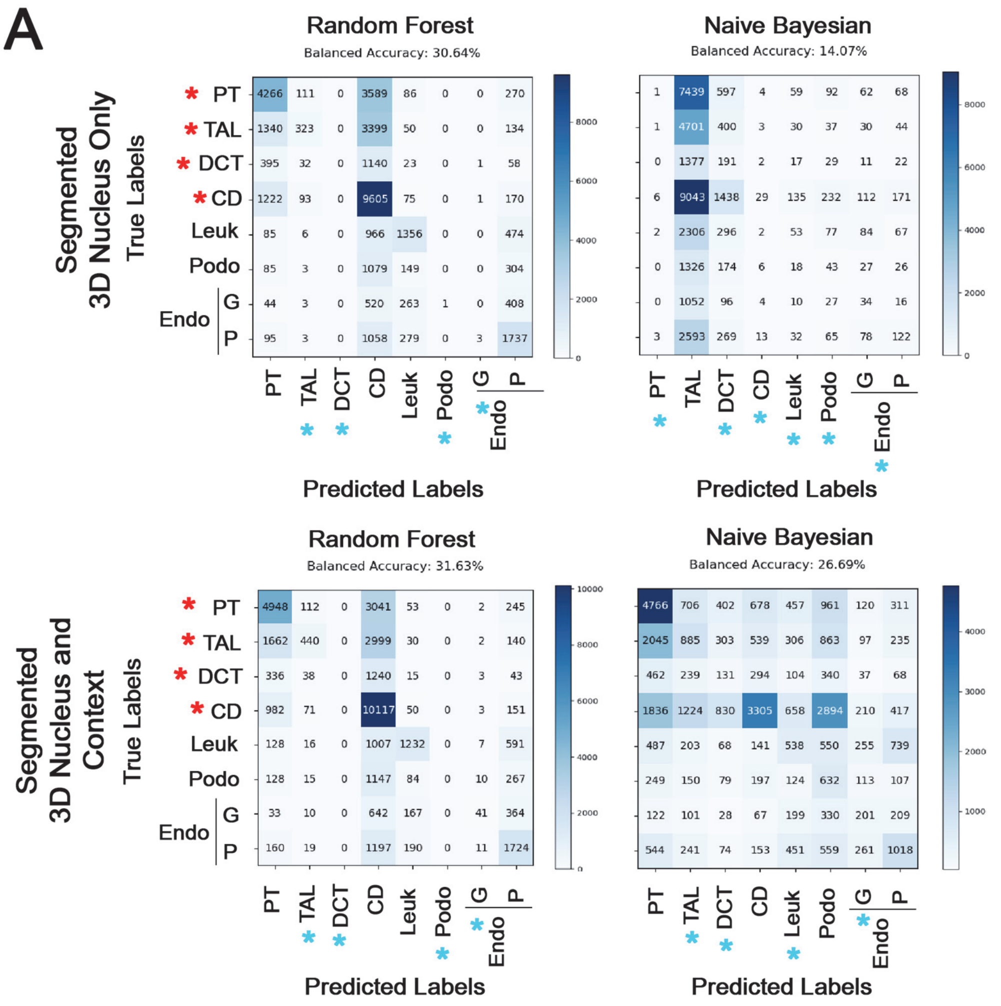


**Figure S2. Cell Classification based on nuclear staining using Random Forest and Naïve Bayesian classifiers.** 2D maximum projections, 3D nuclei or 3D nuclei with context from the 8 class NepNuc datasets were split into training and testing datasets and used to train either a Random Forest or Naïve Bayesian classifier with vectorized images. **A.** Confusion matrices of classifiers for either 3D nuclei or 3D nuclei with context. The eight classes are epithelial cells from the proximal tubule(PT), thick ascending limb (TAL), distal convoluted tubule (DCT), and collecting duct (CD), leukocytes (Leuk), podocytes (Podo) and endothelial cells (Endo) that are either in glomeruli (G) or in the peritubular (P) space. Asterisks indicate error within the epithelial cells (red asterisks), and labels where the classifiers performed poorly (blue asterisks).

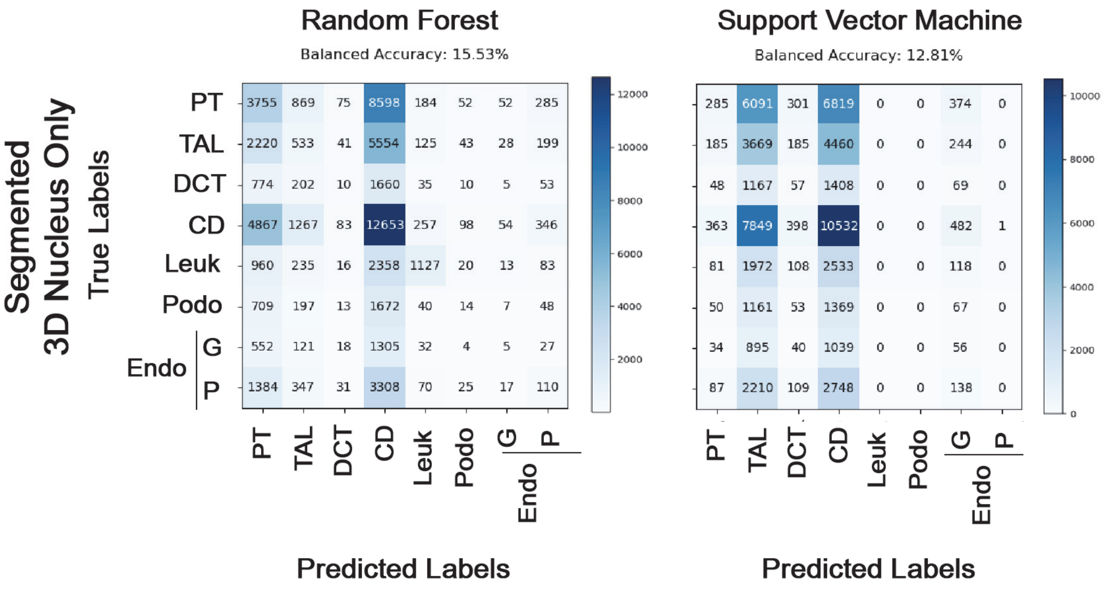


| Random Forest | PT | TAL | DCT | CD | Leuk | Podo | Endo G | Endo P |
|---|---|---|---|---|---|---|---|---|
| PT | 3755 | 869 | 75 | 8598 | 184 | 52 | 52 | 285 |
| TAL | 2220 | 533 | 41 | 5554 | 125 | 43 | 28 | 199 |
| DCT | 774 | 202 | 10 | 1660 | 35 | 10 | 5 | 53 |
| CD | 4867 | 1267 | 83 | 12653 | 257 | 98 | 54 | 346 |
| Leuk | 960 | 235 | 16 | 2358 | 1127 | 20 | 13 | 83 |
| Podo | 709 | 197 | 13 | 1672 | 40 | 14 | 7 | 48 |
| Endo G | 552 | 121 | 18 | 1305 | 32 | 4 | 5 | 27 |
| Endo P | 1384 | 347 | 31 | 3308 | 70 | 25 | 17 | 110 |

| Support Vector Machine | PT | TAL | DCT | CD | Leuk | Podo | Endo G | Endo P |
|---|---|---|---|---|---|---|---|---|
| PT | 285 | 6091 | 301 | 6819 | 0 | 0 | 374 | 0 |
| TAL | 185 | 3669 | 185 | 4460 | 0 | 0 | 244 | 0 |
| DCT | 48 | 1167 | 57 | 1408 | 0 | 0 | 69 | 0 |
| CD | 363 | 7849 | 398 | 10532 | 0 | 0 | 482 | 1 |
| Leuk | 81 | 1972 | 108 | 2533 | 0 | 0 | 118 | 0 |
| Podo | 50 | 1161 | 53 | 1369 | 0 | 0 | 67 | 0 |
| Endo G | 34 | 895 | 40 | 1039 | 0 | 0 | 56 | 0 |
| Endo P | 87 | 2210 | 109 | 2748 | 0 | 0 | 138 | 0 |

**Figure S3. Spherical harmonic-based classification of nuclei.** Triangle meshes of 3D nuclei were generated from the original voxel data, vertices converted to polar coordinates and spherical harmonics were calculated with a least-squares inversion and at most three coefficients. A rotational-invariant frequency spectrum was generated from the modeled spherical harmonic and used as a feature for classification with either **A**, Random Forest or **B**, Support Vector Machine. The classifiers performed poorly across labels.

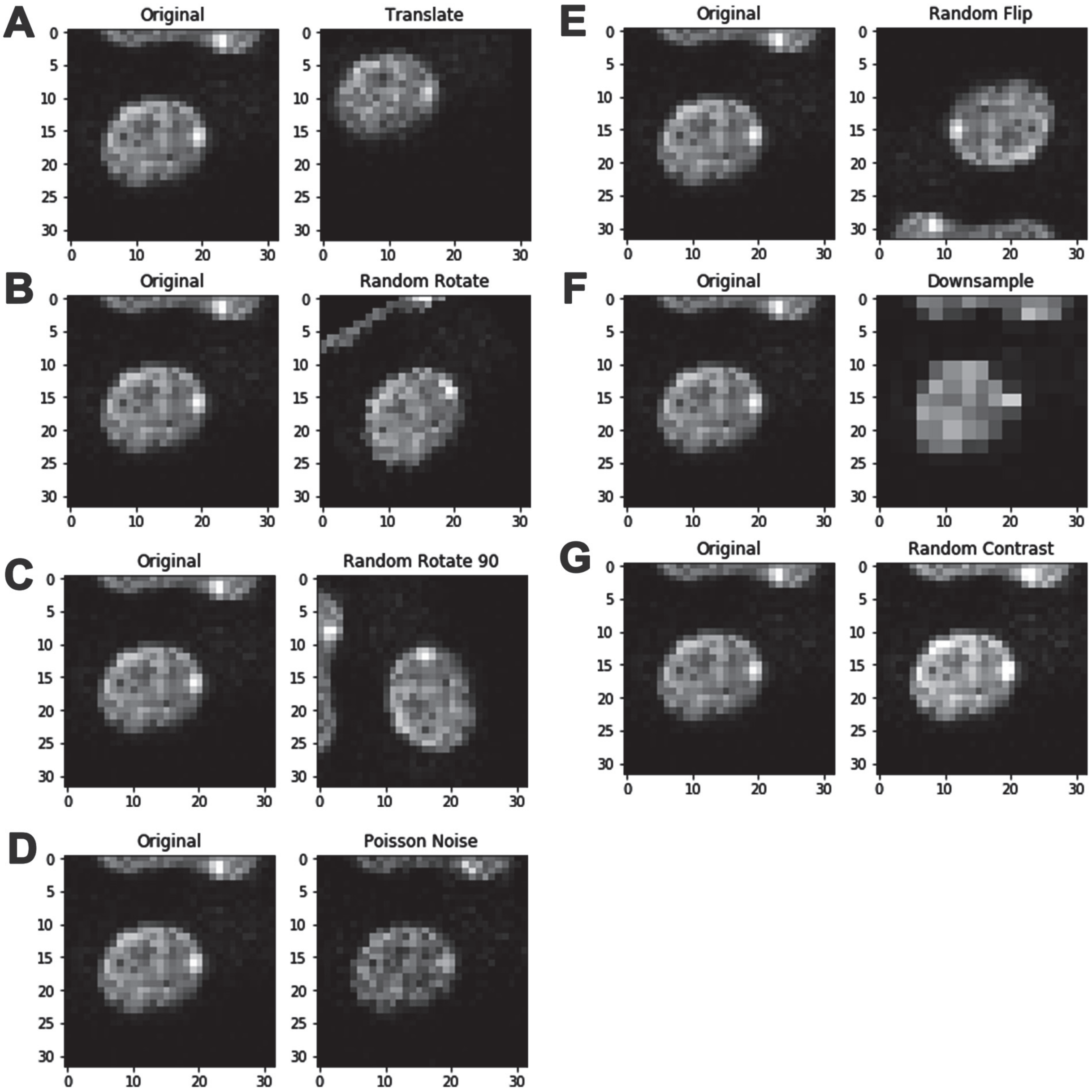


**Figure S4. Augmentation examples on slices from a single volume.** Each augmentation had a 30% chance of being applied to any image. Poisson noise was add using based on a mixing factor, alpha, between the original image and added noise. Contrast was added by multiplying the original image by a contrast factor between 0.8 and 1.2. The image was then clipped to 8-bit depth (0 to 255).

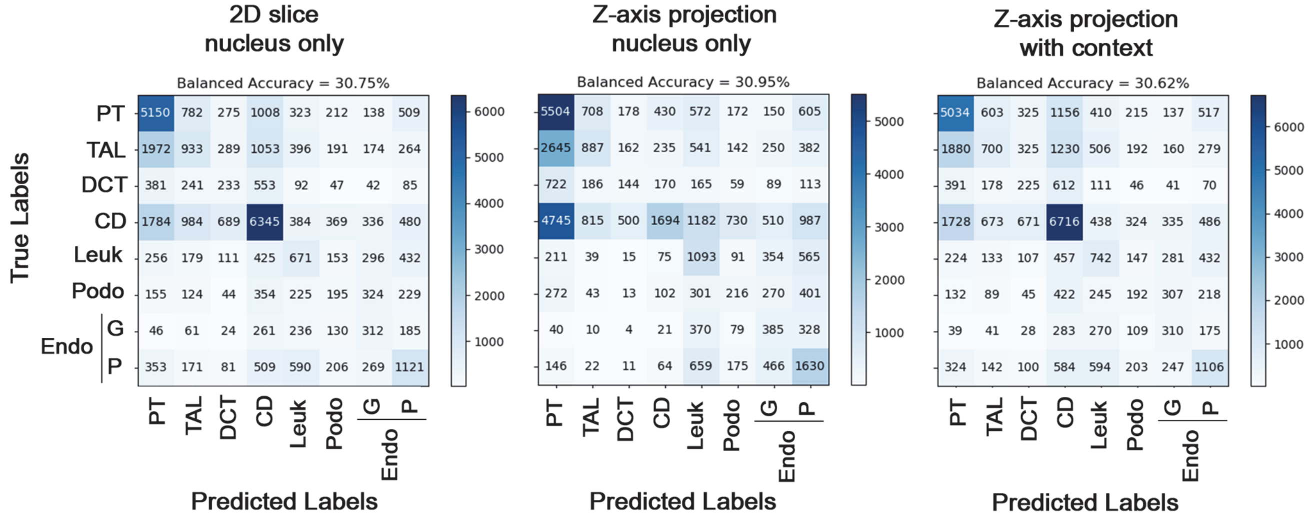


**Figure S5. Transfer learning with ResNet-31.** Ground truth images of nuclei either 2D single slice images or 2D z-axis maximum projections of nuclei with or without surrounding nuclei were split into training and testing datasets and the training datasets were used to train ResNet-31. The testing datasets were classified, and accuracy and confusion matrices were generated. ResNet-31 with 2D or z-projections performs poorly in discriminating between the labels.
